# A genomic perspective across Earth’s microbiomes reveals that genome size in Archaea and Bacteria is linked to ecosystem type and trophic strategy

**DOI:** 10.1101/2021.01.18.427069

**Authors:** Alejandro Rodríguez-Gijón, Julia K. Nuy, Maliheh Mehrshad, Moritz Buck, Frederik Schulz, Tanja Woyke, Sarahi L. Garcia

## Abstract

Our view of genome size in Archaea and Bacteria has remained skewed as the data used to paint its picture has been dominated by genomes of microorganisms that can be cultivated under laboratory settings. However, the continuous effort to catalog the genetic make-up of Earth’s microbiomes specifically propelled by recent extensive work on uncultivated microorganisms, provides a unique opportunity to revise our perspective on genome size distribution. Capitalizing on a recently released extensive catalog of tens of thousands of metagenome-assembled genomes, we provide a comprehensive overview of genome size distributions. We observe that the known phylogenetic diversity of environmental microorganisms possesses significantly smaller genomes than the collection of laboratory isolated microorganisms. Aquatic microorganisms average 3.1 Mb, host-associated microbial genomes average 3.0 Mb, terrestrial microorganism average 3.7 Mb and isolated microorganisms average 4.3 Mb. While the environment where the microorganisms live can certainly be linked to genome size, in some cases, evolutionary phylogenetic history can be a stronger predictor. Moreover, ecological strategies such as auxotrophies have a direct impact on genome size. To better understand the ecological drivers of genome size, we expand on the known and the overlooked factors that influence genome size in different environments, phylogenetic groups and trophic strategies.

## Introduction

Genomes are dynamic databases that encode the machinery behind the evolution and adaptation of living organisms to environmental settings. In brief, a genome encompasses all genetic material present in one organism and includes both its genes and its non-coding DNA. Genome size is largely a function of expansion and contraction by the gain or loss of DNA fragments. Genomes of extant organisms are the result of a long evolutionary history. In eukaryotes, an organism’s complexity is not directly proportional to its genome size, which can have variations over 64 000-fold (1, 2). However, the genome size ranges in Archaea and Bacteria are smaller and the genomes are information-dense (3). The known genome sizes range from 112 kb in *Candidatus* Nasuia deltocephalinicola (4) to 16.04 Mb in *Minicystis rosea* (5). While subject to genetic drift, prokaryotic and eukaryotic genomes have diverging constraints on size. Bacteria exhibit a mutational bias that deletes superfluous sequences, whereas Eukaryotes are biased toward large insertions (6). In Archaea and Bacteria, evolutionary studies have revealed extremely rapid and highly variable flux of genes (7) with drift promoting genome reduction (8).

Genome size dynamics and evolution in Archaea and Bacteria are quite complex and have been studied by many researchers who each focused on different taxonomic lineages or different ecological or evolutionary backgrounds (8–14). The evolutionary forces driving genome size are debated in many excellent reviews (9, 10, 12, 13, 15). However, in this review, we want to focus on the genome size of Archaea and Bacteria seen from an ecological perspective. As microbial researchers, how do we define what is a small or a big genome? Perhaps, researchers working on model organisms such as *Escherichia coli* with a genome size of ∼5 Mb (16) would define ’big’ or ’small’ very differently to researchers working on soil-dwelling bacteria with a genome size of 16 Mb (5), the abundant *Prochlorococcus* with a genome size of ∼2 Mb (17), or bacterial endosymbionts of insects that may have genomes merely larger than 100 kb (4). The recently published expanded databases of environmental archaeal and bacterial genomes (18, 19) allow us to revisit and acquire a complete understanding of genome size distribution across different environments with a higher resolution. This review offers an overview of the distribution of estimated genome sizes of all known archaeal and bacterial phyla across different environments. We found that while several archaeal phyla and two bacterial phyla with consistently smaller genome sizes (< 2 Mb, Figure 1B), 76.3% of representative archaeal and bacterial genomes recovered through genome-resolved metagenomics present estimated genome sizes below 4 Mb.

**Figure 1.**
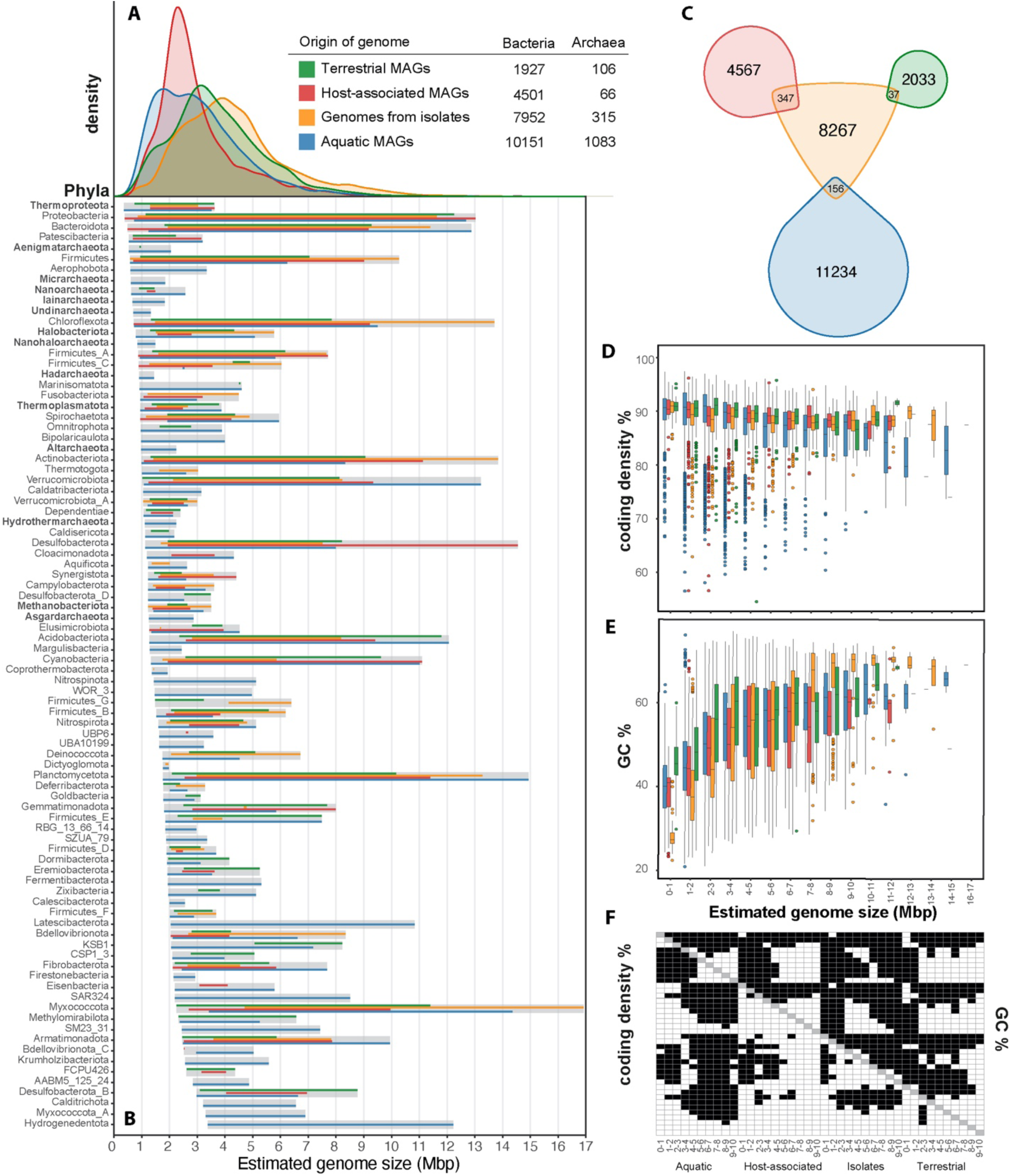
Overview of the genome size distribution across Earth’s microbiomes. Genome size distribution of Archaea and Bacteria [A] from different environmental sources and across different archaeal and bacterial phyla [B] are shown for a total of 26,101 representative genomes. Isolate genomes were gathered from GTDB (release95) and environmental MAGs were gathered from GEMs (18) and stratfreshDB (19). We use one representative genome per mOTU (defined by 95% ANI) from the union of GEMs catalog and stratfreshDB in the plots. From the GTDB database, we selected one representative isolate genome per species cluster that was circumscribed based on the ANI (>=95%) and alignment fraction ((AF) >65%) between genomes (20). To construct the figures, we plotted the estimated genome sizes, which were calculated based on the genome assembly size and completeness estimation provided. Venn diagram of the intersection between the representative environmental MAGs and the representative isolate genomes [C]. The intersection was calculated using FastANI (66) and was determined with a threshold of 95%. The coding density [D] and GC content (%) [E] are shown for the archaeal and bacterial MAGs across different environments and isolates. Pair-wise t-test was performed in all variables of panel E and F and shown in [F], where white is significant (p<0.05) and black is not significant (p>0.05). In panel B, we only included phyla with more than five genomes.

## Extant genome size distribution in the environment

The current state of environmental sequencing, assembly, and binning technologies allows us to review and renew our view of archaeal and bacterial genome size distribution on Earth (18–20). To minimize representation biases (21), from the ∼64 500 environmental metagenome-assembled genomes (MAGs), we included one representative per mOTU, defined by 95% average nucleotide identity (ANI), from the GEMs and the stratfreshDB MAGs resulting in ∼18 000 MAGs. We complemented these data by adding ∼8 000 species (or mOTUs) cluster representatives from >90% complete genomes of isolates from GTDB (Figure 1A). We found 540 mOTUs with representatives in both the environmental MAGs and the isolate genomes (Figure 1C). This would mean that about 3% of the extracted MAGs from the environment have a cultivated representative.

Furthermore, using completeness estimates from CheckM (22), we compared the genome size distribution of all MAGs versus genomes from isolates. Isolates have an average genome size of 4.3 Mb which is significantly larger than that of MAGs (t-test p<0.0001), both when comparing Archaea and Bacteria combined and separately. Although the ecosystem classification we have chosen to display is coarse and might contain countless niches, it still allowed us to see trends for genome sizes. Aquatic MAGs average 3.1 Mb, host-associated MAGs average 3.0 Mb, and terrestrial MAGs average 3.7 Mb (Figure 1A). It is known that MAG assembly might discriminate against ribosomal RNAs, transfer RNAs, mobile element functions and genes of unknown function (23, 24), and also that completeness estimations can be underestimated for streamlined genomes (25). For the 540 mOTUs with MAGs and isolate genomes (Figure 1C), we found that MAGs were estimated on average 3.7% smaller than isolate genomes (Figure S1). This suggests that there might be only a small bias in metagenome assembly and binning of these environmental genomes. On its own, it would not account for the genome size difference between all isolate representatives and all MAGs.

A reason for the difference in genome size between isolates and microorganisms living in different ecosystems might be related to the fact that traditional isolation techniques select for rare microorganisms (26) and do not capture the entire ecosystem’s diversity (Figure 1C). For example, it is known that current cultivation techniques with rich media bias the cultivation towards copiotrophic and fast-growing microorganisms (27). Moreover, microorganisms in nature do not live in isolation but have coevolved with other microorganisms and might have specific requirements that are hard to meet in batch-culture standard-media isolation techniques (28). Other reasons for biases in cultivation include slow growth of microorganisms (29), host dependency (30), dormancy (31), and microorganisms with very limited metabolic capacity (32) among others. More innovations to culturing the uncultured microbial majority (33) will enable us to bring representatives from the whole genome size spectrum to culture.

Placing archaeal and bacterial genome sizes in phylogenetic trees (Figure 2A and B) shows that the distribution of representative genomes and their estimated sizes varies widely between different phyla and within phyla. Eight phyla in the domain Archaea were reconstructed exclusively from aquatic environments, whereas eight other archaeal phyla were found in multiple ecosystems. There was no significant difference between the genome sizes of those two groups of archaeal phyla (Figure 2C). However, estimated genome sizes in bacterial phyla were significantly larger than those in archaeal phyla. Moreover, genera from phyla with genome sizes below 3 Mb, such as Halobacteriota, Thermoproteota and Patescibacteria, do not show genome size variation in different ecosystems (Figure 2D, 2E, 2I). Nevertheless, genera from smaller genome size phyla are significantly smaller than genera with more genome size variation in any environment (Figure 2K-2N). For phyla spanning genome sizes above 3 Mb, the genome sizes in aquatic or host-associated genera are smaller than terrestrial or non-specific environments (Figure 2F, 2G, 2H, 2J). We observe that while the microorganisms’ environment can certainly be linked to genome size, evolutionary phylogenetic history can be a stronger predictor in phyla where genome sizes are mostly below 3 Mb.

**Figure 2.**
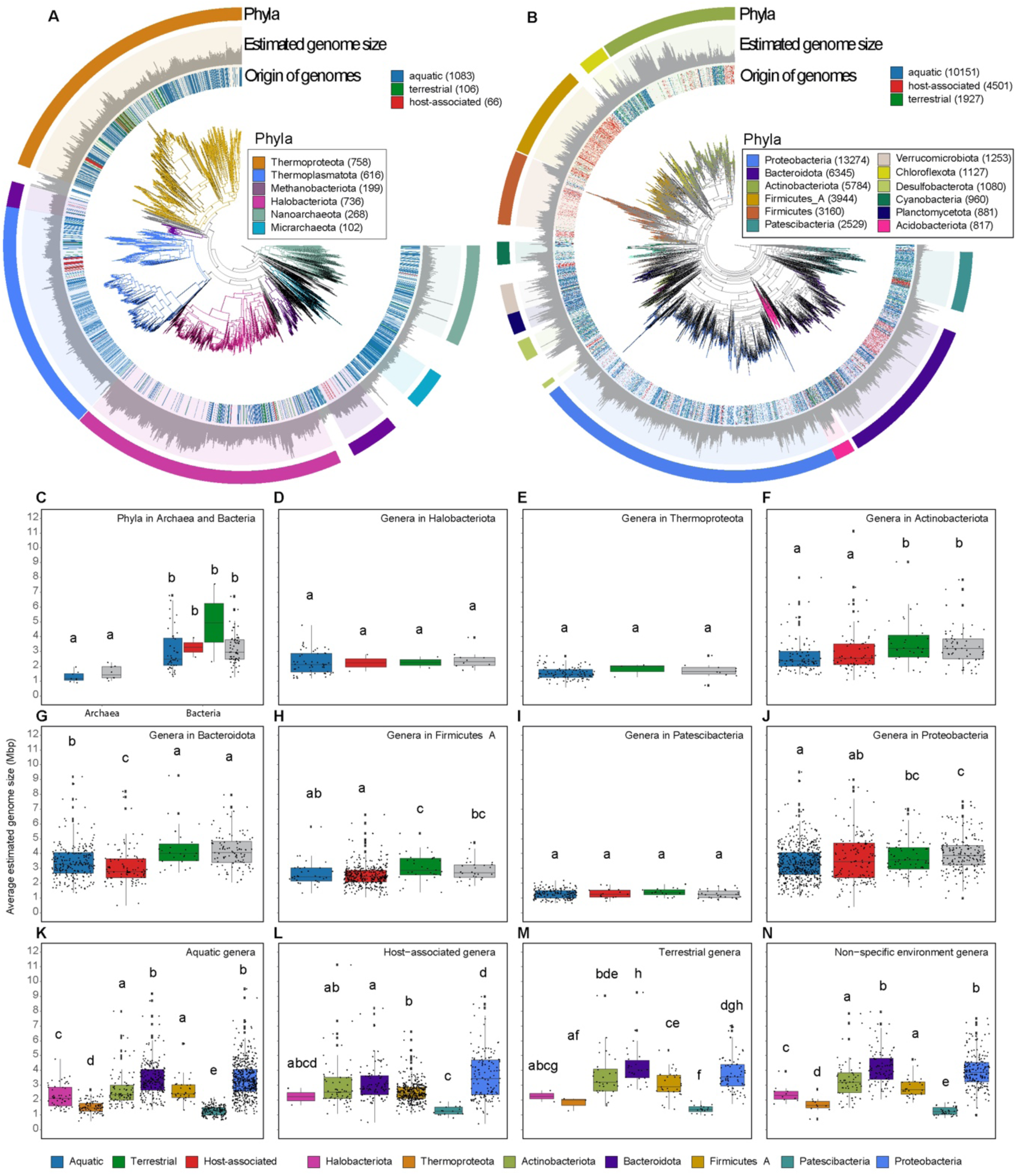
Phylogenetic trees of archaeal [A] and bacterial [B] representative genomes show variation in genome size between and within phyla. The trees were constructed using GTDB-tk using *de novo* workflow using aligned concatenated set of 122 and 120 single copy marker proteins for Archaea and Bacteria, respectively (67). Moreover, in this mode, GTDB-tk adds 1672 and 30238 backbone genomes for Archaea and Bacteria, respectively. Estimated genome size is in scale from 0 Mbp to 6 Mpb or 14 Mbp for Archaea or Bacteria respectively and it shows the distribution of larger and smaller genomes sizes are non-monophyletic. In the tree, the origin of the environmental genomes is labeled: aquatic, terrestrial and host-associated (same MAGs as Figure 1). Highlighted phyla with more representative genomes are color-coded. Boxplots show the average estimated genome size per phyla within Archaea and Bacteria [C] domain. The average estimated size per genus within Halobacteriota [D], Thermoproteota [E], Actinobacteriota [F], Bacteroidota [G], Firmicutes A [H], Patescibacteria [I], Proteobacteria [J]. The presence of the genus is marked as non-specific when there are MAGs in it whose origin is not the same environment. The average estimated size per genus extracted from aquatic environments [K], host-associated ecosystems [L], terrestrial environments [M], or non-specific environments [N]. Letters in boxplot panels are the result of non-parametric tests, Wilcoxon and/or Kruskal-Wallis. Different letters show significant differences p<0.05

Lumping microorganisms together by the three biome categories is not optimal since each biome contains innumerable niches, each of which will have different selective pressures on the genome size. An example is clearly shown in a study (34) in which it is observed that Archaea and Bacteria sampled from different parts of the human body have differences in genome size. Low metadata resolution and clustering of all genomes into three main environments might be a reason why we see a range of genome sizes in the genera of different environments (Figure 2). With more precise metadata and higher sampling resolution of micro-areas, it might be possible to better identify the ecological drivers of genome sizes in the different niches in the environment. But for now, we will discuss the known and the overlooked ecological drivers of genome sizes.

## Impact of ecosystem and trophic strategy on genome size

Terrestrial ecosystems harbor immense microbial diversity (35). Yet, the most up-to-date data compilation provided here shows only 2033 MAGs from terrestrial environments (Figure 1C) with an average genome size of 3.7 Mb (Figure 1A). The sub-ecosystems considered in this view are soil and deep subsurface, among others (Figure S2). While the terrestrial microorganism’s genome size is the biggest of the three ecosystem categories in this review, they are smaller than expected based on previous metagenomic predictions, which placed the genome size of soil bacteria at 4.74 Mb (36). Trends of larger genome sizes in soil have been hypothesized to be related to scarcity and high diversity of nutrients, fluctuating environment combined with little penalty for the slow growth rate (11, 37, 38). Although terrestrial environments are physically structured, they are generally characterized by two to three orders of magnitude greater variations (in temperature and currents) than marine environments (39). *In silico* studies predict that large genome sizes could result from higher environmental variability (40). A recent example showed that isolates of terrestrial Cyanobacteria have genomes on the larger size scale (6.0-8.0. Mb) that are enriched in genes involved in regulatory, transport and motility functions (37). These functional categories enable thriving in a fluctuating environment and high nutrient diversity. Despite these general trends showing larger genome sizes in terrestrial environments, it is worth noting that the diversity captured in the GEMs survey is probably a small fraction of the total terrestrial microbial diversity. It is, for example, also known that streamlined microorganisms such as Patescibacteria (Fig 1B) and ‘*Candidatus* Udaeobacter copiosus’ (Verrucomicrobiota) are abundant in soils (41). We predict that the view on genome size distribution in terrestrial ecosystems will be more complete with more sequencing, assembly, binning and novel isolation efforts.

In host-associated microbiomes, genetic drift, deletion biases and low populations sizes drive the reduction of genomes. In these environments, microorganisms are shaped in their ecological and evolutionary history by the differing levels of intimacy they might have with their host. For example, within the Chlamydiaceae family, some lineages have evolved intracellular associations with eukaryotes (42, 43). These intracellular Chlamydiaceae have lost many genes when comparing them to their common ancestor Chlamydiia (class) that lives in the environment (44). Moreover, host-associated bacterial genomes show a variation in size depending on the type of host (plant, animal, etc.) and the type of association they have with the host, such as endosymbiotic, ectobiotic, or epibiotic (Table S1). Generally, microorganisms associated with Arthropoda (45), humans (46) and other mammals show smaller genomes sizes, whereas protist- and plant-associated bacteria present larger genomes (47) (Figure S2). In fact, *in silico* studies of Alphaproteobacteria show massive genome expansions diversifying plant-associated Rhizobiales and extreme gene losses in the ancestor of the intracellular lineages Rickettsia, Wolbachia, Bartonella and Brucella that are animal- and human-associated (48). Although host-associated microorganisms are widely known for their reduced genomes, the characteristics of host-associated MAGs show coding densities of ∼91% for genomes between 0 and 2 Mb (Figure 1D).

Small genomes exhibit either strong dependency on other community members or have specific nutrient requirements. Two diverging views on genome reduction have emerged. On the one hand, genetic drift is more pronounced in species that have a small effective population size, such as host-associated endosymbiotic microorganisms. These microorganisms might thrive because hosts provide energy or nutrients. On the other hand, streamlining is the process of gene loss through selection and it is mainly observed in free-living microorganisms with high effective population sizes. Some of the most numerically abundant and streamlined microorganisms known to date, such as Pelagibacter (class Alphaproteobacteria) (10), *Prochlorococcus* (phylum Cyanobacteria) (17) Thermoproteota (49) and Patescibacteria (50), are commonly found in aquatic niches. Paradoxically, even though these microorganisms are free-living, their small genomes increase their nutritional connectivity to other individuals (10). Free-living aquatic microorganisms have been used as exemplary streamlining cases in which many have gone through community adaptive selections and gene loss (51). Their gene loss goes so far that they become auxotrophic, meaning they cannot biosynthesize essential metabolites. One strategy to overcome their required nutritional needs is to thrive in functional cohorts (52). As opposed to prototrophic lifestyle, auxotrophic lifestyle is reflected by smaller genome sizes (25, 41, 53, 54) (Table S1). In our freshwater survey (19), from a total of 887 mOTUs with at least 3 MAGs each, only 61 had the metabolic potential to biosynthesize vitamin B12 *de novo* (Figure 3B). Moreover, genomes of Vitamin B12 synthesizers were on average 1 Mb larger than Vitamin B12 auxotrophs (Figure 3C). An exciting avenue for future studies includes understanding how prevalent auxotrophies are for the entire spectrum of metabolites (amino acids, nucleotides, fatty acids, vitamins, etc.) in different microbial communities and how those auxotrophies are linked with genome size.

**Figure 3.**
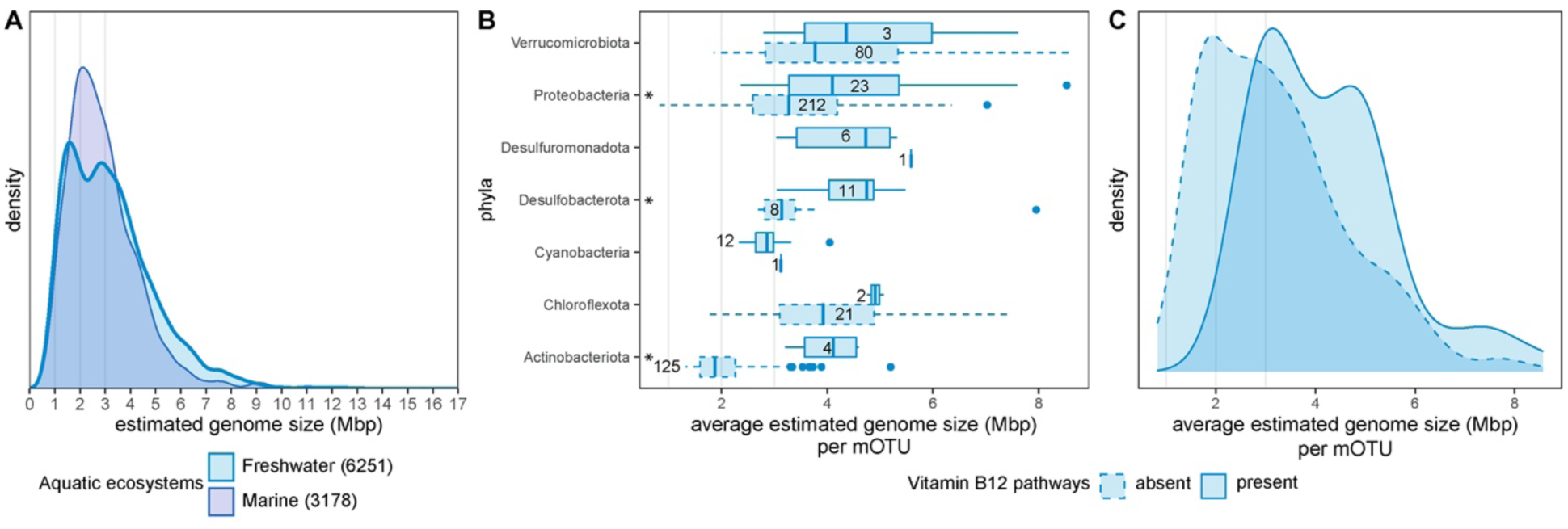
Genome size distribution of estimated genome sizes of representative MAGs recovered from marine and freshwater ecosystems [A]. The number of MAGs in each sub-ecosystem is in parenthesis. Average genome size distribution of mOTUs within each phylum from the stratfreshDB with present and absent vitamin B12 biosynthesis capacity [B]. Phyla with no Vitamin B12 synthesizer were excluded. The presence and absence of oxic and anoxic Vitamin B12 capacities were analyzed by assessing the completeness of KEGG modules. A module was considered present (solid boxplots) when its completeness was >= 81.67 %, which is the average completeness of mOTUs in the stratfreshDB. A module was considered absent when none of the genes assigned to any of the vitamin B12 modules was detected in the mOTU (dashed boxplots). Wilcoxon signed-rank tests were performed to test for significant differences (p-value <= 0.05) of average estimated genome sizes in mOTUs with absent and present Vitamin B12 pathways within a phylum. Significant differences are marked with an asterisk. The number of mOTUs is shown in or next to the corresponding boxplot. Distribution of average estimated genome sizes per mOTU with and without VitaminB12 biosynthesis capacity [C].

In this review, the largest fraction of MAGs is recovered from aquatic environments. The two main sub-ecosystems show that freshwater MAGs (average 3.2 Mb) bimodal genome size distribution is significantly different (p<0.0001) from the unimodal marine genome size distribution (average 2.9 Mb) (Figure 3A). Although the potential to synthesize Vitamin B12 might partly explain the bimodal distribution of genome sizes in freshwaters (Figure 3C), more research is necessary to fully understand the trophic strategies behind the bimodal freshwater genome size distribution. Moreover, when comparing freshwater and marine environments, the most obvious difference is salinity followed by nutrient concentration. Further exploring the impact of differing levels of salinity on genome size is an interesting research prospect. In general, aquatic environments are vertically structured by gradients of light penetration, temperature, oxygen, and nutrient. Moreover, microorganisms might experience a microscale spatial and nutrient structure due to the presence of heterogeneous particles. These aquatic structures are drivers of the genetic repertoire of aquatic microorganisms. Metagenomic sequencing reported the increase of genome sizes for Archaea and Bacteria with increasing depths (55). Temperature may be as important; for example, a study based on twenty-one Thermoproteota and Euryarchaeota fosmids (Euryarchaetoa is now reclassified into Methanobacteriota, Halobacteriota and Nanohaloarchaeota) showed high rates of gene gains through HGT to adapt to cold and deep marine environments (56). One other driver we want to point out in aquatic environments is light which decreases with depth. Photosynthetic bacteria such as *Prochlorococcus* spp. are well-differentiated into a high-light adapted ecotype with smaller genome sizes (average 1.6 Mb) and a low-light-adapted ecotype with a slightly bigger genome size (average 1.9 Mb) (57) (Table S1). Limitation of nutrients such as nitrogen (58) might also be one of the central factors determining genomic properties (59). Nitrogen fixation is a complex process that requires several genes (60) and most nitrogen-fixing marine cyanobacteria have the largest genomes (61).

Diversity and quantity of nutrients might be two understudied factors that drive ecology and genome size evolution. A recent example shows that polysaccharide xylan triggers microcolonies, whereas monosaccharide xylose promotes solitary growth in Caulobacter (62). This is a striking example of how nutrient complexity can foster diverse niches for well-studied cells such as Caulobacter with genome size 4 Mb. We believe that to fully understand the link between genome size and nutritional requirements of diverse environmental microorganisms, we need to systematically explore the ∼90% of molecules/metabolites still unknown (63–65). The wide nutrient complexity in the environment might prompt microorganisms to shape their genome. Their genome content will allow them either to feed or not on a variety of nutrients and might leave them either depending or not on other microorganisms. Metagenomics combined with metabolomics will provide an understanding of the genome size of microorganisms and their nutritional and trophic strategy.

## Conclusion

This review offers an overview where genomes obtained from environmental samples show to be smaller than those obtained from laboratory isolates. This is not mainly because isolates and MAGs from the same species differed in size but because cultivation methods bias the sampling of environmental microbiome towards obtaining copiotrophs, fast growers, and more metabolically independent microorganisms. Moreover, we find the distribution of genome sizes across the phylogenetic tree of Archaea and Bacteria can be linked to the environment where the microorganisms live. In some cases, phylogenetic history can be a stronger predictor of genome size than the environment. Finally, we review the ecological factors causing the varying sizes of genomes in different ecosystems. Soils might have the microorganisms with the bigger estimated genome sizes due to higher fluctuations in the environment. Host-associations might shape genomes sizes differentially based on the type of host and level of intimacy between the microorganisms and the host. Genomes in aquatic environments might be shaped by vertical stratification in nutrients and light penetration and particle distribution. Moreover, different trophic strategies such as auxotrophies might be connected to smaller genome sizes. We expect that as the microbial ecology field keeps moving forward with sequencing, bioinformatics, chemical analysis, and novel cultivation techniques, we will get a deeper resolution on physicochemical, metabolic, spatial, and biological drivers of archaeal and bacterial genome sizes.

## Acknowledgments

We are grateful to John Paul Balmonte and Sergio Tusso for helpful discussions. ARG thanks Fede Berckx for technical advice. This work was supported by SciLifeLab and Kungl. Vetenskapsakademiens stiftelser grant CR2019-0060. The computations and data handling were enabled by resources in the project SNIC 2021/6-99 and SNIC 2021/5-133 provided by the Swedish National Infrastructure for Computing (SNIC) at UPPMAX, partially funded by the Swedish Research Council through grant agreement no. 2018-05973. The work conducted by the U.S. Department of Energy Joint Genome Institute, an Office of Science User Facility, made use of the National Energy Research Scientific Computing Center and was supported under Contract No. DE-AC02-05CH11231.

## Contributions

SLG, ARG and JN conceptualized the literature and data review idea. JN, MB and FS gathered the data. ARG and JN performed data analysis. All authors did literature searches. SLG, ARG and MM drafted the first manuscript. All authors contributed to the writing and editing of the manuscript.

---

**Figure S1.**
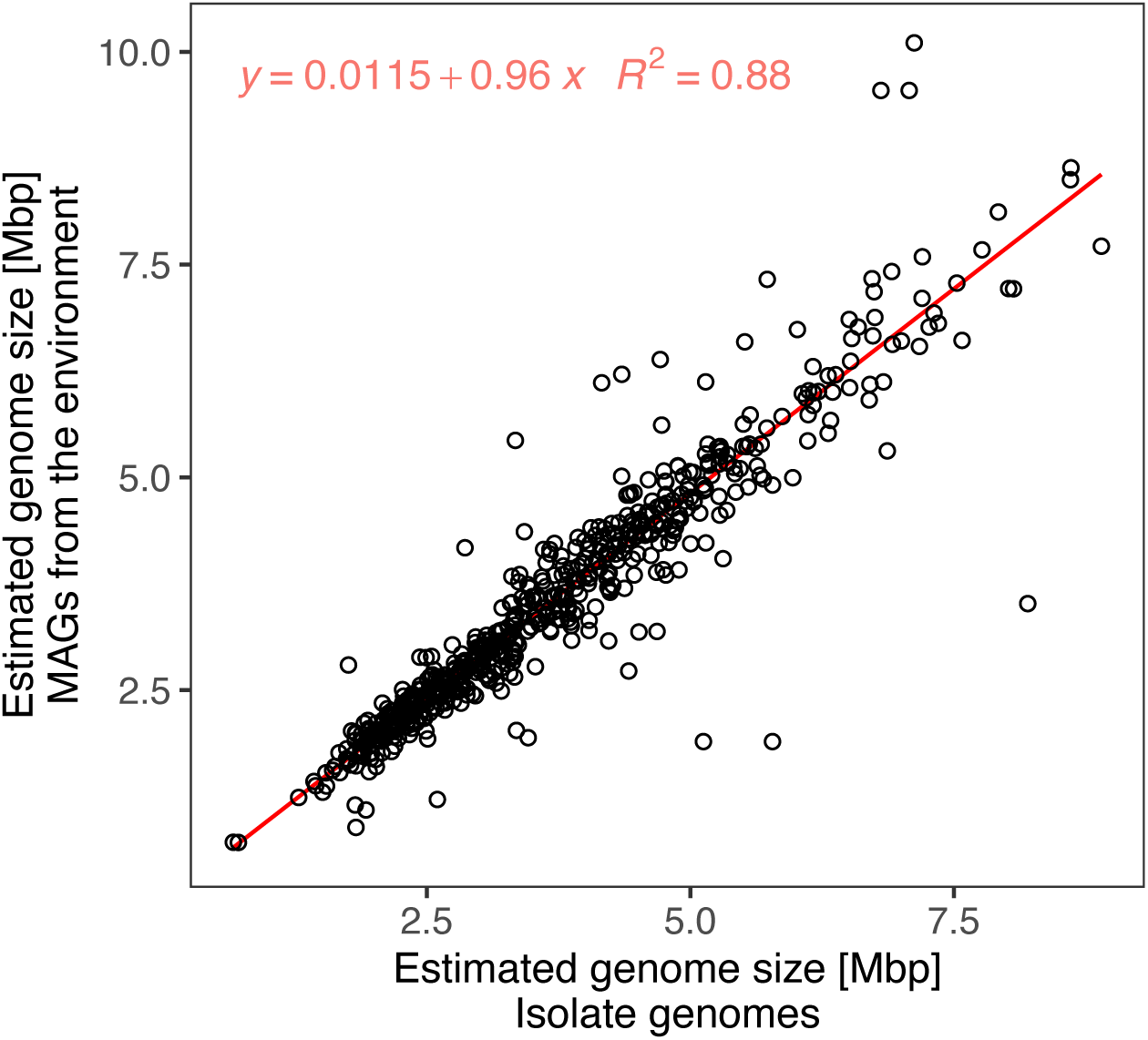
Comparison of conspecific MAGs and isolate genomes. In total, 17834 representative MAGs from environments were clustered with 8267 reference genomes from isolates into mOTUs at 95% ANI. Only 560 MAGs formed clusters with 556 isolate genomes resulting in 540 mOTUs. Each point in the plot represents a MAG/mOTU pair assigned to a single mOTU. The x-axis indicates the estimated genome size of isolates genomes and the y-axis indicates the estimated genome size of MAGs.

**Figure S2.**
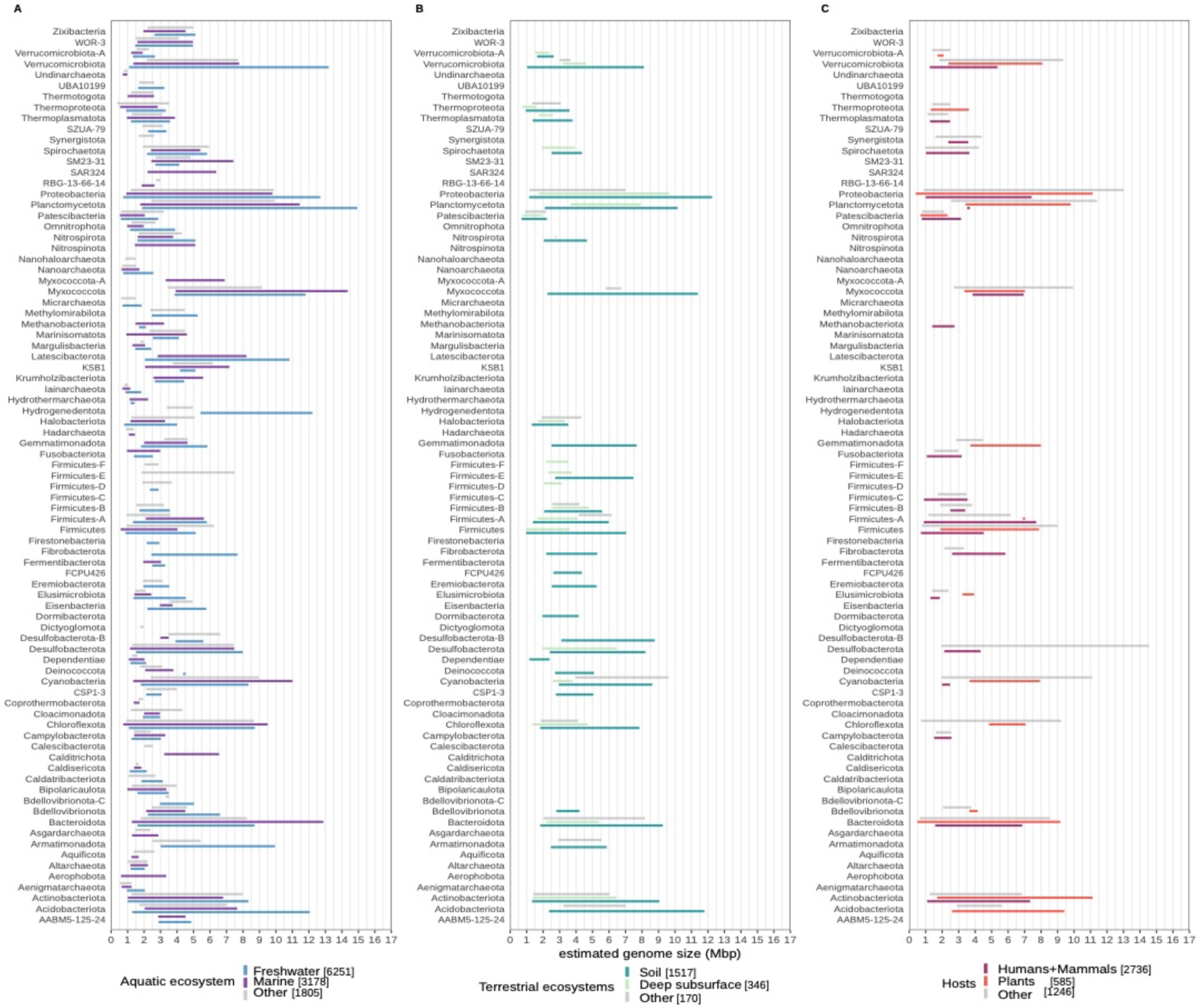
Distribution of estimated genome sizes per phyla in aquatic [A], terrestrial [B], and host-associated ecosystems [C]. In each Panel, the two sub ecosystems are shown from which the most MAGs were recovered, while ’Others’ combine MAGs from less represented sub ecosystems.

**Table S1.**
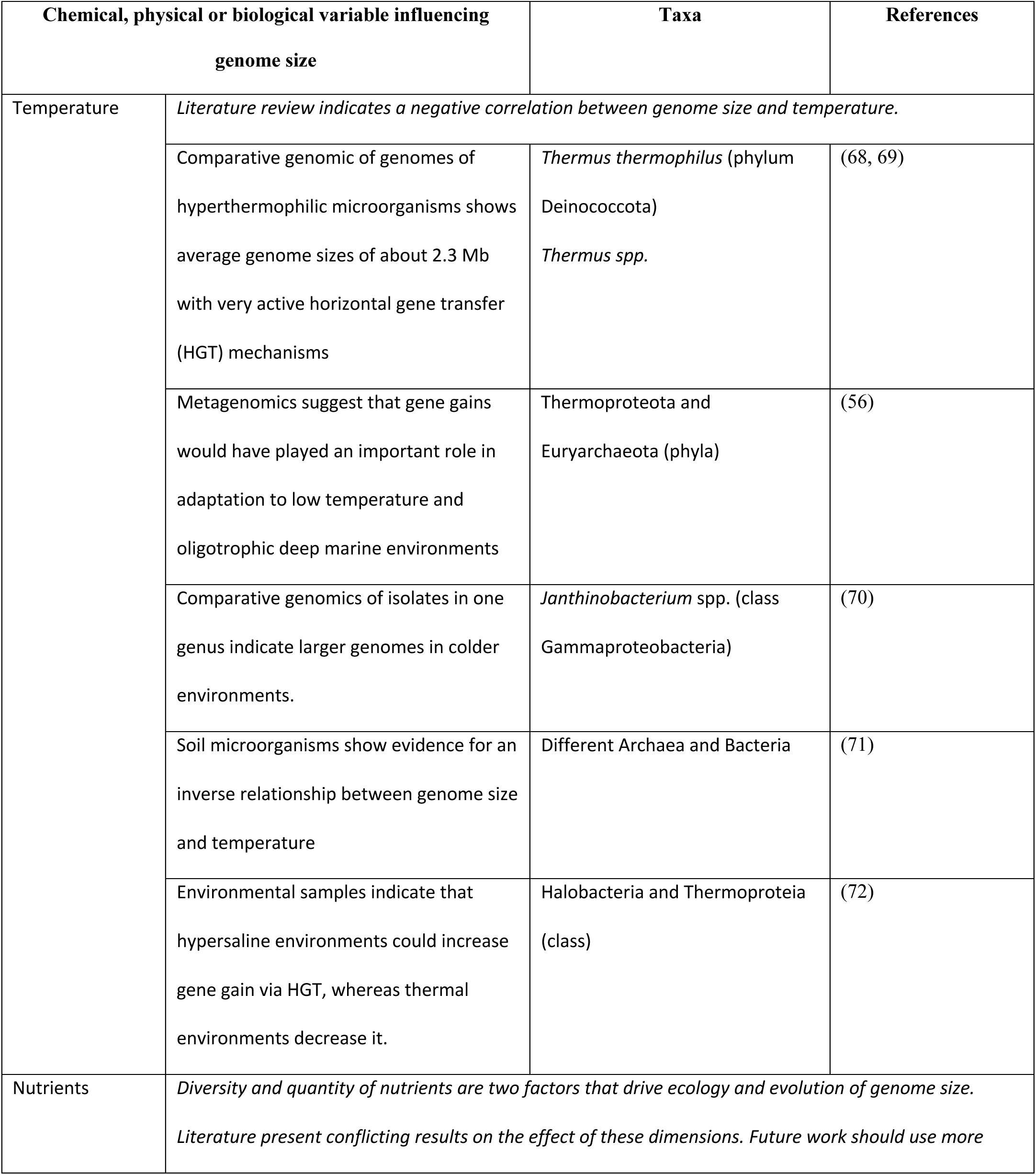
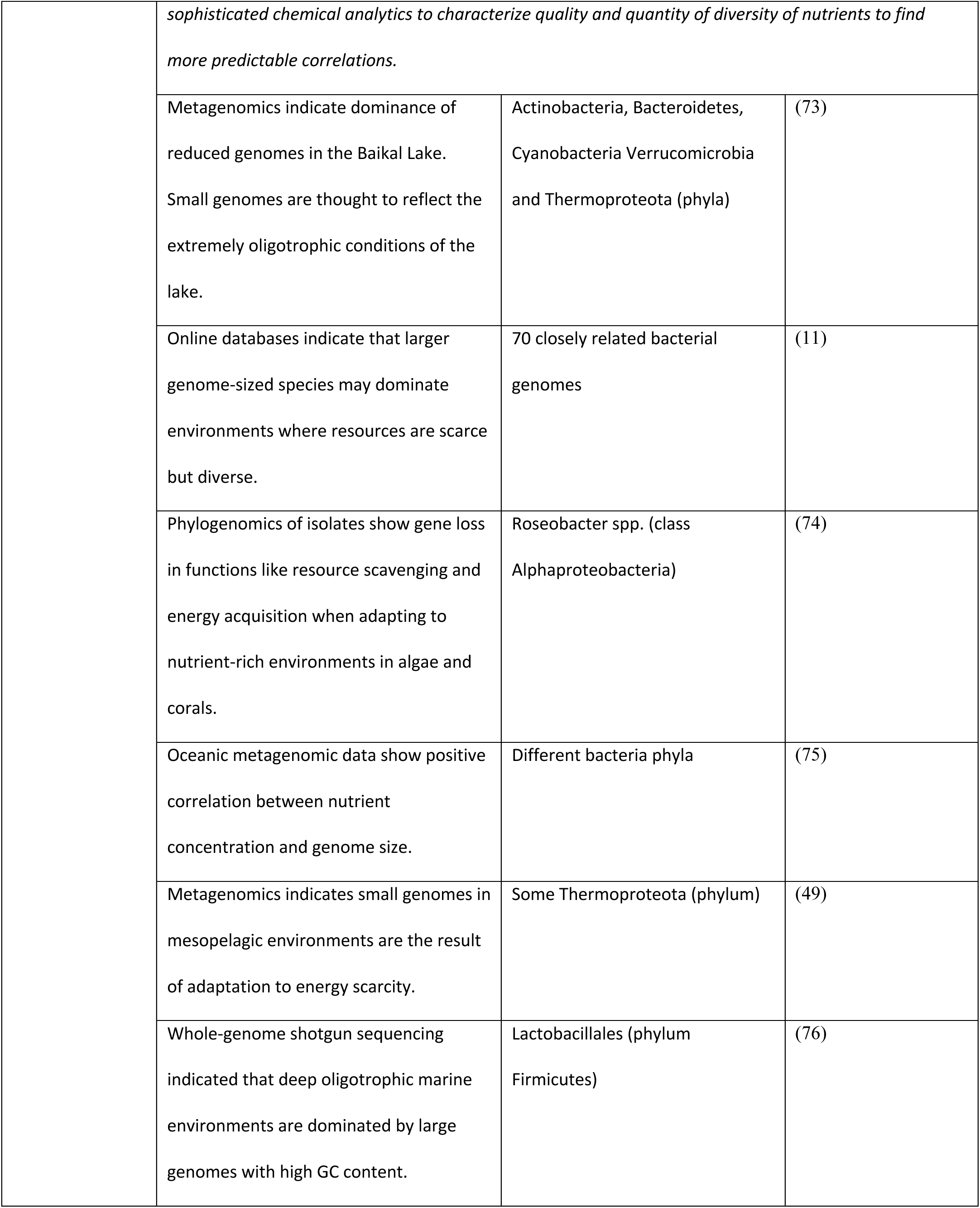
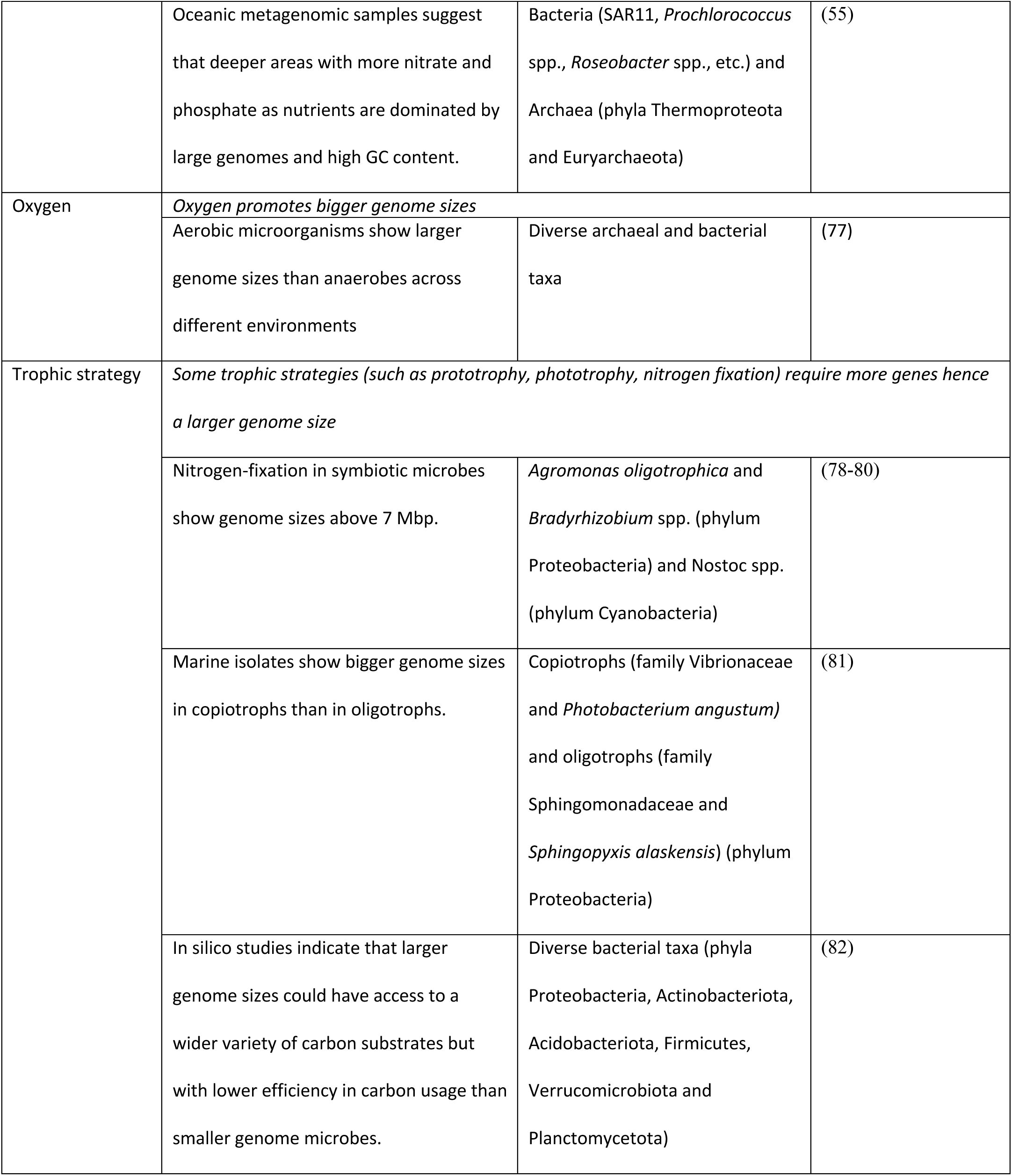
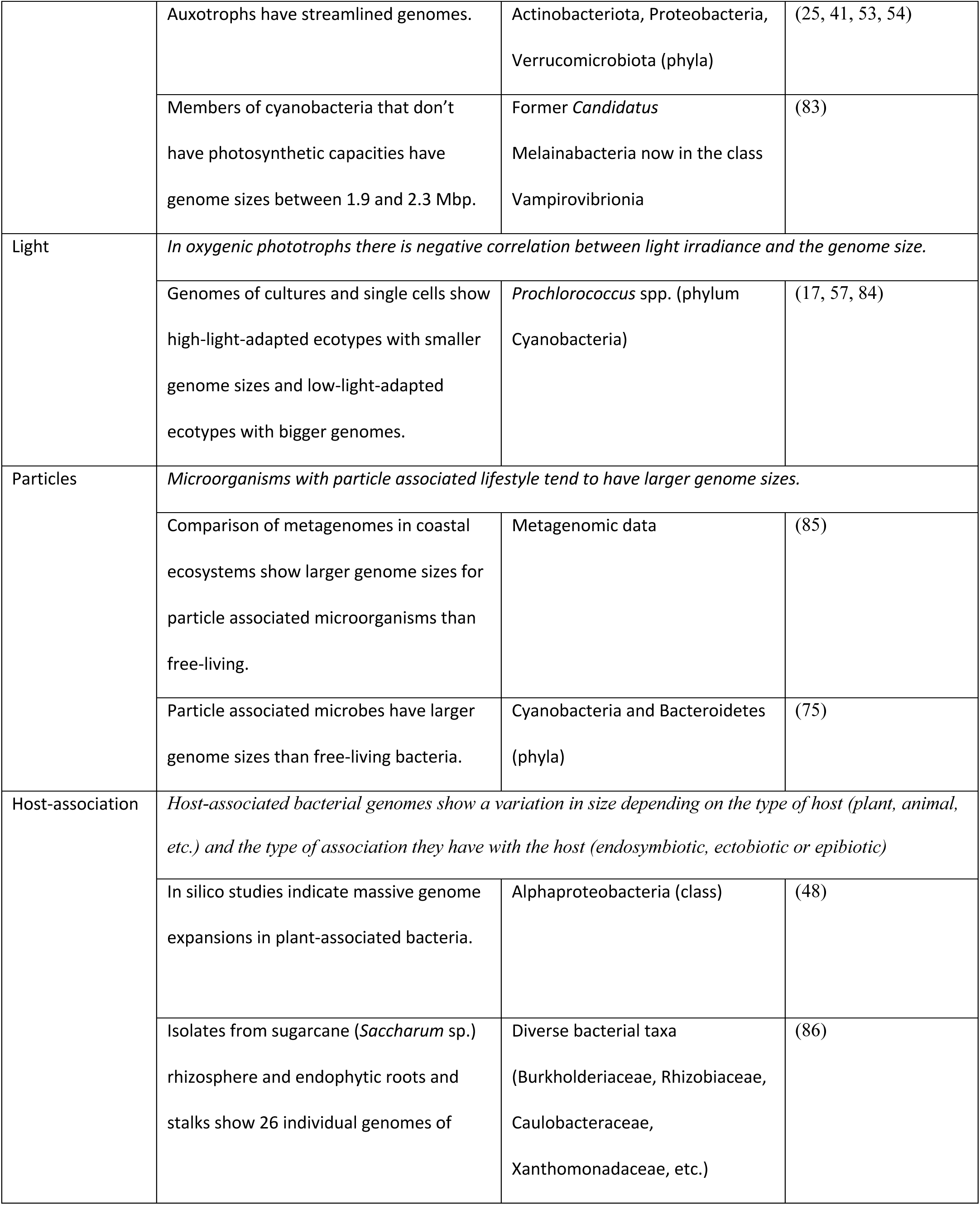
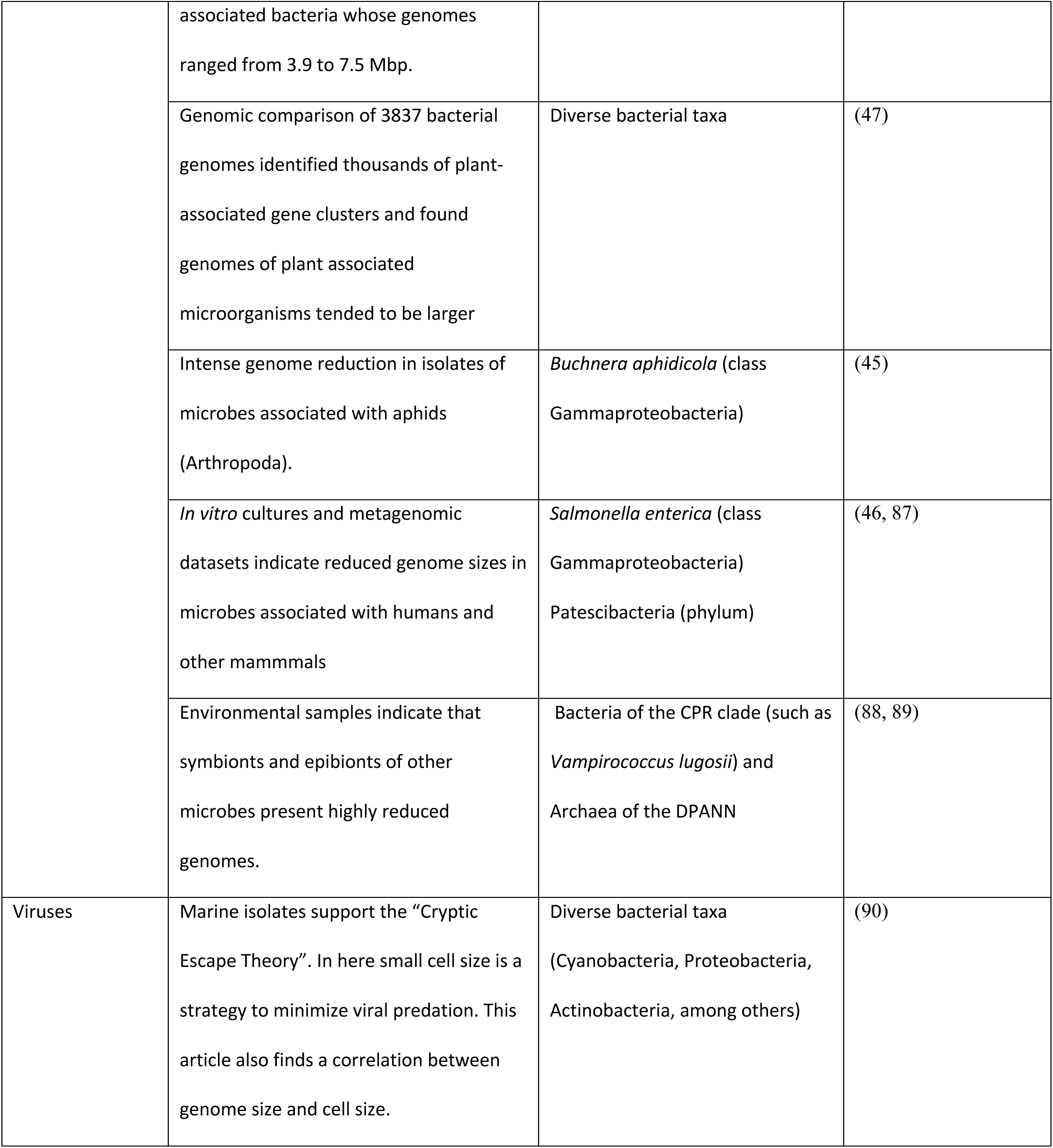
Ecological factors and their correlation to genome size.

## References

1. Gregory TR. 2005. Genome Size Evolution in Animals, p 3–87, The Evolution of the Genome doi:10.1016/b978-012301463-4/50003-6.

2. Pellicer J, Hidalgo O, Dodsworth S, Leitch IJ. 2018. Genome Size Diversity and Its Impact on the Evolution of Land Plants. Genes (Basel) 9.

3. Kirchberger PC, Schmidt ML, Ochman H. 2020. The Ingenuity of Bacterial Genomes. Annu Rev Microbiol 74:815–834.

4. Moran NA, Bennett GM. 2014. The tiniest tiny genomes. Annu Rev Microbiol 68:195–215.

5. Garcia R, Gemperlein K, Muller R. 2014. Minicystis rosea gen. nov., sp. nov., a polyunsaturated fatty acid-rich and steroid-producing soil myxobacterium. Int J Syst Evol Microbiol 64:3733–3742.

6. Bobay LM, Ochman H. 2017. The Evolution of Bacterial Genome Architecture. Front Genet 8:72.

7. Puigbo P, Lobkovsky AE, Kristensen DM, Wolf YI, Koonin EV. 2014. Genomes in turmoil: quantification of genome dynamics in prokaryote supergenomes. BMC Biol 12:66.

8. Kuo CH, Moran NA, Ochman H. 2009. The consequences of genetic drift for bacterial genome complexity. Genome Res 19:1450–4.

9. Batut B, Knibbe C, Marais G, Daubin V. 2014. Reductive genome evolution at both ends of the bacterial population size spectrum. Nat Rev Microbiol 12:841–50.

10. Giovannoni SJ, Cameron Thrash J, Temperton B. 2014. Implications of streamlining theory for microbial ecology. ISME J 8:1553–65.

11. Konstantinidis KT, Tiedje JM. 2004. Trends between gene content and genome size in prokaryotic species with larger genomes. Proc Natl Acad Sci U S A 101:3160–5.

12. Lynch M. 2006. Streamlining and simplification of microbial genome architecture. Annu Rev Microbiol 60:327–49.

13. Wolf YI, Koonin EV. 2013. Genome reduction as the dominant mode of evolution. Bioessays 35:829–37.

14. Bourguignon T, Kinjo Y, Villa-Martin P, Coleman NV, Tang Q, Arab DA, Wang Z, Tokuda G, Hongoh Y, Ohkuma M, Ho SYW, Pigolotti S, Lo N. 2020. Increased Mutation Rate Is Linked to Genome Reduction in Prokaryotes. Curr Biol 30:3848–3855 e4.

15. Mira A, Ochman H, Moran NA. 2001. Deletional bias and the evolution of bacterial genomes. Trends in Genetics 17:589–596.

16. Abram K, Udaondo Z, Bleker C, Wanchai V, Wassenaar TM, Robeson MS, 2nd, Ussery DW. 2021. Mash-based analyses of Escherichia coli genomes reveal 14 distinct phylogroups. Commun Biol 4:117.

17. Rocap G, Larimer FW, Lamerdin J, Malfatti S, Chain P, Ahlgren NA, Arellano A, Coleman M, Hauser L, Hess WR, Johnson ZI, Land M, Lindell D, Post AF, Regala W, Shah M, Shaw SL, Steglich C, Sullivan MB, Ting CS, Tolonen A, Webb EA, Zinser ER, Chisholm SW. 2003. Genome divergence in two Prochlorococcus ecotypes reflects oceanic niche differentiation. Nature 424:1042–7.

18. Nayfach S, Roux S, Seshadri R, Udwary D, Varghese N, Schulz F, Wu D, Paez-Espino D, Chen IM, Huntemann M, Palaniappan K, Ladau J, Mukherjee S, Reddy TBK, Nielsen T, Kirton E, Faria JP, Edirisinghe JN, Henry CS, Jungbluth SP, Chivian D, Dehal P, Wood-Charlson EM, Arkin AP, Tringe SG, Visel A, Consortium IMD, Woyke T, Mouncey NJ, Ivanova NN, Kyrpides NC, Eloe-Fadrosh EA. 2020. A genomic catalog of Earth’s microbiomes. Nat Biotechnol doi:10.1038/s41587-020-0718-6.

19. Buck M, Garcia SL, Fernandez L, Martin G, Martinez-Rodriguez GA, Saarenheimo J, Zopfi J, Bertilsson S, Peura S. 2021. Comprehensive dataset of shotgun metagenomes from oxygen stratified freshwater lakes and ponds. Sci Data 8:131.

20. Parks DH, Chuvochina M, Chaumeil PA, Rinke C, Mussig AJ, Hugenholtz P. 2020. A complete domain-to-species taxonomy for Bacteria and Archaea. Nat Biotechnol 38:1079–1086.

21. Gweon HS, Bailey MJ, Read DS. 2017. Assessment of the bimodality in the distribution of bacterial genome sizes. ISME J 11:821–824.

22. Parks DH, Imelfort M, Skennerton CT, Hugenholtz P, Tyson GW. 2015. CheckM: assessing the quality of microbial genomes recovered from isolates, single cells, and metagenomes. Genome Res doi:10.1101/gr.186072.114.

23. Nelson WC, Tully BJ, Mobberley JM. 2020. Biases in genome reconstruction from metagenomic data. PeerJ 8:e10119.

24. Meziti A, Rodriguez RL, Hatt JK, Pena-Gonzalez A, Levy K, Konstantinidis KT. 2021. The Reliability of Metagenome-Assembled Genomes (MAGs) in Representing Natural Populations: Insights from Comparing MAGs against Isolate Genomes Derived from the Same Fecal Sample. Appl Environ Microbiol 87.

25. Garcia SL, Buck M, McMahon KD, Grossart HP, Eiler A, Warnecke F. 2015. Auxotrophy and intrapopulation complementary in the “interactome’ of a cultivated freshwater model community. Molecular Ecology 24:4449–4459.

26. Shade A, Hogan CS, Klimowicz AK, Linske M, McManus PS, Handelsman J. 2012. Culturing captures members of the soil rare biosphere. Environmental Microbiology 14:2247–2252.

27. Swan BK, Tupper B, Sczyrba A, Lauro FM, Martinez-Garcia M, Gonzalez JM, Luo H, Wright JJ, Landry ZC, Hanson NW, Thompson BP, Poulton NJ, Schwientek P, Acinas SG, Giovannoni SJ, Moran MA, Hallam SJ, Cavicchioli R, Woyke T, Stepanauskas R. 2013. Prevalent genome streamlining and latitudinal divergence of planktonic bacteria in the surface ocean. Proc Natl Acad Sci U S A 110:11463–8.

28. Garcia SL. 2016. Mixed cultures as model communities: hunting for ubiquitous microorganisms, their partners, and interactions. Aquatic Microbial Ecology 77:79–85.

29. Imachi H, Nobu MK, Nakahara N, Morono Y, Ogawara M, Takaki Y, Takano Y, Uematsu K, Ikuta T, Ito M, Matsui Y, Miyazaki M, Murata K, Saito Y, Sakai S, Song C, Tasumi E, Yamanaka Y, Yamaguchi T, Kamagata Y, Tamaki H, Takai K. 2020. Isolation of an archaeon at the prokaryote-eukaryote interface. Nature 577:519–525.

30. Cross KL, Campbell JH, Balachandran M, Campbell AG, Cooper SJ, Griffen A, Heaton M, Joshi S, Klingeman D, Leys E, Yang Z, Parks JM, Podar M. 2019. Targeted isolation and cultivation of uncultivated bacteria by reverse genomics. Nat Biotechnol 37:1314–1321.

31. Hoehler TM, Jorgensen BB. 2013. Microbial life under extreme energy limitation. Nat Rev Microbiol 11:83–94.

32. Figueroa-Gonzalez PA, Bornemann TLV, Adam PS, Plewka J, Revesz F, von Hagen CA, Tancsics A, Probst AJ. 2020. Saccharibacteria as Organic Carbon Sinks in Hydrocarbon-Fueled Communities. Front Microbiol 11:587782.

33. Lewis WH, Tahon G, Geesink P, Sousa DZ, Ettema TJG. 2020. Innovations to culturing the uncultured microbial majority. Nat Rev Microbiol doi:10.1038/s41579-020-00458-8.

34. Nayfach S, Pollard KS. 2015. Average genome size estimation improves comparative metagenomics and sheds light on the functional ecology of the human microbiome. Genome Biol 16:51.

35. Delgado-Baquerizo M, Oliverio AM, Brewer TE, Benavent-Gonzalez A, Eldridge DJ, Bardgett RD, Maestre FT, Singh BK, Fierer N. 2018. A global atlas of the dominant bacteria found in soil. Science 359:320–325.

36. Raes J, Korbel J, Lercher M, von Mering C, Bork P. 2007. Prediction of effective genome size in metagenomic samples. Genome Biology 8:R10.

37. Chen MY, Teng WK, Zhao L, Hu CX, Zhou YK, Han BP, Song LR, Shu WS. 2021. Comparative genomics reveals insights into cyanobacterial evolution and habitat adaptation. ISME J 15:211–227.

38. Cobo-Simon M, Tamames J. 2017. Relating genomic characteristics to environmental preferences and ubiquity in different microbial taxa. BMC Genomics 18:499.

39. Steele JH, Brink KH, Scott BE. 2019. Comparison of marine and terrestrial ecosystems: suggestions of an evolutionary perspective influenced by environmental variation. ICES Journal of Marine Science 76:50–59.

40. Bentkowski P, Van Oosterhout C, Mock T. 2015. A Model of Genome Size Evolution for Prokaryotes in Stable and Fluctuating Environments. Genome Biol Evol 7:2344–51.

41. Brewer TE, Handley KM, Carini P, Gilbert JA, Fierer N. 2016. Genome reduction in an abundant and ubiquitous soil bacterium ‘Candidatus Udaeobacter copiosus’. Nature Microbiology 2:16198.

42. Toft C, Andersson SG. 2010. Evolutionary microbial genomics: insights into bacterial host adaptation. Nat Rev Genet 11:465–75.

43. Collingro A, Kostlbacher S, Horn M. 2020. Chlamydiae in the Environment. Trends Microbiol 28:877–888.

44. Dharamshi JE, Tamarit D, Eme L, Stairs CW, Martijn J, Homa F, Jorgensen SL, Spang A, Ettema TJG. 2020. Marine Sediments Illuminate Chlamydiae Diversity and Evolution. Curr Biol 30:1032–1048 e7.

45. Tamas I, Klasson L, Canback B, Naslund AK, Eriksson AS, Wernegreen JJ, Sandstrom JP, Moran NA, Andersson SG. 2002. 50 million years of genomic stasis in endosymbiotic bacteria. Science 296:2376–9.

46. McLean JS, Bor B, Kerns KA, Liu Q, To TT, Solden L, Hendrickson EL, Wrighton K, Shi W, He X. 2020. Acquisition and Adaptation of Ultra-small Parasitic Reduced Genome Bacteria to Mammalian Hosts. Cell Rep 32:107939.

47. Levy A, Salas Gonzalez I, Mittelviefhaus M, Clingenpeel S, Herrera Paredes S, Miao J, Wang K, Devescovi G, Stillman K, Monteiro F, Rangel Alvarez B, Lundberg DS, Lu TY, Lebeis S, Jin Z, McDonald M, Klein AP, Feltcher ME, Rio TG, Grant SR, Doty SL, Ley RE, Zhao B, Venturi V, Pelletier DA, Vorholt JA, Tringe SG, Woyke T, Dangl JL. 2017. Genomic features of bacterial adaptation to plants. Nat Genet 50:138–150.

48. Boussau B, Karlberg EO, Frank AC, Legault BA, Andersson SG. 2004. Computational inference of scenarios for alpha-proteobacterial genome evolution. Proc Natl Acad Sci U S A 101:9722–7.

49. Aylward FO, Santoro AE. 2020. Heterotrophic Thaumarchaea with Small Genomes Are Widespread in the Dark Ocean. mSystems 5.

50. Tian R, Ning D, He Z, Zhang P, Spencer SJ, Gao S, Shi W, Wu L, Zhang Y, Yang Y, Adams BG, Rocha AM, Detienne BL, Lowe KA, Joyner DC, Klingeman DM, Arkin AP, Fields MW, Hazen TC, Stahl DA, Alm EJ, Zhou J. 2020. Small and mighty: adaptation of superphylum Patescibacteria to groundwater environment drives their genome simplicity. Microbiome 8:51.

51. Morris JJ, Lenski RE, Zinser ER. 2012. The Black Queen Hypothesis: evolution of dependencies through adaptive gene loss. MBio 3.

52. Mondav R, Bertilsson S, Buck M, Langenheder S, Lindstrom ES, Garcia SL. 2020. Streamlined and Abundant Bacterioplankton Thrive in Functional Cohorts. mSystems 5.

53. Kang I, Kim S, Islam MR, Cho J-C. 2017. The first complete genome sequences of the acI lineage, the most abundant freshwater Actinobacteria, obtained by whole-genome-amplification of dilution-to-extinction cultures. Scientific Reports 7.

54. Grote J, Thrash JC, Huggett MJ, Landry ZC, Carini P, Giovannoni SJ, Rappe MS. 2012. Streamlining and Core Genome Conservation among Highly Divergent Members of the SAR11 Clade. Mbio 3.

55. Mende DR, Bryant JA, Aylward FO, Eppley JM, Nielsen T, Karl DM, DeLong EF. 2017. Environmental drivers of a microbial genomic transition zone in the ocean’s interior. Nat Microbiol 2:1367–1373.

56. Brochier-Armanet C, Deschamps P, Lopez-Garcia P, Zivanovic Y, Rodriguez-Valera F, Moreira D. 2011. Complete-fosmid and fosmid-end sequences reveal frequent horizontal gene transfers in marine uncultured planktonic archaea. ISME J 5:1291–302.

57. Berube PM, Biller SJ, Hackl T, Hogle SL, Satinsky BM, Becker JW, Braakman R, Collins SB, Kelly L, Berta-Thompson J, Coe A, Bergauer K, Bouman HA, Browning TJ, De Corte D, Hassler C, Hulata Y, Jacquot JE, Maas EW, Reinthaler T, Sintes E, Yokokawa T, Lindell D, Stepanauskas R, Chisholm SW. 2018. Single cell genomes of Prochlorococcus, Synechococcus, and sympatric microbes from diverse marine environments. Sci Data 5:180154.

58. Elser JJ, Bracken ME, Cleland EE, Gruner DS, Harpole WS, Hillebrand H, Ngai JT, Seabloom EW, Shurin JB, Smith JE. 2007. Global analysis of nitrogen and phosphorus limitation of primary producers in freshwater, marine and terrestrial ecosystems. Ecol Lett 10:1135–42.

59. Grzymski JJ, Dussaq AM. 2012. The significance of nitrogen cost minimization in proteomes of marine microorganisms. ISME J 6:71–80.

60. Franche C, Lindström K, Elmerich C. 2008. Nitrogen-fixing bacteria associated with leguminous and non-leguminous plants. Plant and Soil 321:35–59.

61. Bergman B, Sandh G, Lin S, Larsson J, Carpenter EJ. 2013. Trichodesmium--a widespread marine cyanobacterium with unusual nitrogen fixation properties. FEMS Microbiol Rev 37:286–302.

62. D’Souza GG, Povolo VR, Keegstra JM, Stocker R, Ackermann M. 2021. Nutrient complexity triggers transitions between solitary and colonial growth in bacterial populations. ISME J doi:10.1038/s41396-021-00953-7.

63. Wienhausen G, Noriega-Ortega BE, Niggemann J, Dittmar T, Simon M. 2017. The Exometabolome of Two Model Strains of the Roseobacter Group: A Marketplace of Microbial Metabolites. Front Microbiol 8:1985.

64. Hawkes JA, Patriarca C, Sjöberg PJR, Tranvik LJ, Bergquist J. 2018. Extreme isomeric complexity of dissolved organic matter found across aquatic environments. Limnology and Oceanography Letters 3:21–30.

65. Patriarca C, Sedano-Núñez VT, Garcia SL, Bergquist J, Bertilsson S, Sjöberg PJR, Tranvik LJ, Hawkes JA. 2020. Character and environmental lability of cyanobacteria-derived dissolved organic matter. Limnology and Oceanography 66:496–509.

66. Jain C, Rodriguez-R LM, Phillippy AM, Konstantinidis KT, Aluru S. 2018. High throughput ANI analysis of 90K prokaryotic genomes reveals clear species boundaries. Nature Communications 9.

67. Chaumeil PA, Mussig AJ, Hugenholtz P, Parks DH. 2020. GTDB-Tk: a toolkit to classify genomes with the Genome Taxonomy Database. Bioinformatics 36:1925–1927.

68. Borges KM, Bergquist PL. 1993. Genomic restriction map of the extremely thermophilic bacterium Thermus thermophilus HB8. J Bacteriol 175:103–10.

69. Blesa A, Averhoff B, Berenguer J. 2018. Horizontal Gene Transfer in Thermus spp. Curr Issues Mol Biol 29:23–36.

70. Dieser M, Smith HJ, Ramaraj T, Foreman CM. 2019. Janthinobacterium CG23_2: Comparative Genome Analysis Reveals Enhanced Environmental Sensing and Transcriptional Regulation for Adaptation to Life in an Antarctic Supraglacial Stream. Microorganisms 7.

71. Sorensen JW, Dunivin TK, Tobin TC, Shade A. 2019. Ecological selection for small microbial genomes along a temperate-to-thermal soil gradient. Nat Microbiol 4:55–61.

72. Rhodes ME, Spear JR, Oren A, House CH. 2011. Differences in lateral gene transfer in hypersaline versus thermal environments. BMC Evol Biol 11:199.

73. Cabello-Yeves PJ, Zemskaya TI, Rosselli R, Coutinho FH, Zakharenko AS, Blinov VV, Rodriguez-Valera F. 2018. Genomes of Novel Microbial Lineages Assembled from the Sub-Ice Waters of Lake Baikal. Appl Environ Microbiol 84.

74. Chu X, Li S, Wang S, Luo D, Luo H. 2020. Gene loss through pseudogenization contributes to the ecological diversification of a generalist Roseobacter lineage. ISME J 15:489–502.

75. Allen LZ, Allen EE, Badger JH, McCrow JP, Paulsen IT, Elbourne LD, Thiagarajan M, Rusch DB, Nealson KH, Williamson SJ, Venter JC, Allen AE. 2012. Influence of nutrients and currents on the genomic composition of microbes across an upwelling mosaic. ISME J 6:1403–14.

76. Makarova K, Slesarev A, Wolf Y, Sorokin A, Mirkin B, Koonin E, Pavlov A, Pavlova N, Karamychev V, Polouchine N, Shakhova V, Grigoriev I, Lou Y, Rohksar D, Lucas S, Huang K, Goodstein DM, Hawkins T, Plengvidhya V, Welker D, Hughes J, Goh Y, Benson A, Baldwin K, Lee JH, Diaz-Muniz I, Dosti B, Smeianov V, Wechter W, Barabote R, Lorca G, Altermann E, Barrangou R, Ganesan B, Xie Y, Rawsthorne H, Tamir D, Parker C, Breidt F, Broadbent J, Hutkins R, O’Sullivan D, Steele J, Unlu G, Saier M, Klaenhammer T, Richardson P, Kozyavkin S, Weimer B, Mills D. 2006. Comparative genomics of the lactic acid bacteria. Proc Natl Acad Sci U S A 103:15611–6.

77. Nielsen DA, Fierer N, Geoghegan JL, Gillings MR, Gumerov V, Madin JS, Moore L, Paulsen IT, Reddy TBK, Tetu SG, Westoby M. 2021. Aerobic bacteria and archaea tend to have larger and more versatile genomes. Oikos 130:501–511.

78. Okubo T, Fukushima S, Itakura M, Oshima K, Longtonglang A, Teaumroong N, Mitsui H, Hattori M, Hattori R, Hattori T, Minamisawa K. 2013. Genome analysis suggests that the soil oligotrophic bacterium Agromonas oligotrophica (Bradyrhizobium oligotrophicum) is a nitrogen-fixing symbiont of Aeschynomene indica. Appl Environ Microbiol 79:2542–51.

79. Bromfield ESP, Cloutier S, Nguyen HDT. 2019. Description and complete genome sequence of Bradyrhizobium amphicarpaeae sp. nov., harbouring photosystem and nitrogen-fixation genes. Int J Syst Evol Microbiol 69:2841–2848.

80. Gagunashvili AN, Andresson OS. 2018. Distinctive characters of Nostoc genomes in cyanolichens. BMC Genomics 19:434.

81. Lauro FM, McDougald D, Thomas T, Williams TJ, Egan S, Rice S, DeMaere MZ, Ting L, Ertan H, Johnson J, Ferriera S, Lapidus A, Anderson I, Kyrpides N, Munk AC, Detter C, Han CS, Brown MV, Robb FT, Kjelleberg S, Cavicchioli R. 2009. The genomic basis of trophic strategy in marine bacteria. Proc Natl Acad Sci U S A 106:15527–33.

82. Saifuddin M, Bhatnagar JM, Segre D, Finzi AC. 2019. Microbial carbon use efficiency predicted from genome-scale metabolic models. Nat Commun 10:3568.

83. Di Rienzi SC, Sharon I, Wrighton KC, Koren O, Hug LA, Thomas BC, Goodrich JK, Bell JT, Spector TD, Banfield JF, Ley RE. 2013. The human gut and groundwater harbor non-photosynthetic bacteria belonging to a new candidate phylum sibling to Cyanobacteria. Elife 2:e01102.

84. Dufresne A, Salanoubat M, Partensky F, Artiguenave F, Axmann IM, Barbe V, Duprat S, Galperin MY, Koonin EV, Le Gall F, Makarova KS, Ostrowski M, Oztas S, Robert C, Rogozin IB, Scanlan DJ, Tandeau de Marsac N, Weissenbach J, Wincker P, Wolf YI, Hess WR. 2003. Genome sequence of the cyanobacterium Prochlorococcus marinus SS120, a nearly minimal oxyphototrophic genome. Proc Natl Acad Sci U S A 100:10020–5.

85. Smith MW, Zeigler Allen L, Allen AE, Herfort L, Simon HM. 2013. Contrasting genomic properties of free-living and particle-attached microbial assemblages within a coastal ecosystem. Front Microbiol 4:120.

86. de Souza RSC, Armanhi JSL, Damasceno NB, Imperial J, Arruda P. 2019. Genome Sequences of a Plant Beneficial Synthetic Bacterial Community Reveal Genetic Features for Successful Plant Colonization. Front Microbiol 10:1779.

87. Nilsson AI, Koskiniemi S, Eriksson S, Kugelberg E, Hinton JC, Andersson DI. 2005. Bacterial genome size reduction by experimental evolution. Proc Natl Acad Sci U S A 102:12112–6.

88. Moreira D, Le Guyader H, Philippe H. 2000. The origin of red algae and the evolution of chloroplasts. Nature 405:69–72.

89. Castelle CJ, Banfield JF. 2018. Major New Microbial Groups Expand Diversity and Alter our Understanding of the Tree of Life. Cell 172:1181–1197.

90. Yooseph S, Nealson KH, Rusch DB, McCrow JP, Dupont CL, Kim M, Johnson J, Montgomery R, Ferriera S, Beeson K, Williamson SJ, Tovchigrechko A, Allen AE, Zeigler LA, Sutton G, Eisenstadt E, Rogers YH, Friedman R, Frazier M, Venter JC. 2010. Genomic and functional adaptation in surface ocean planktonic prokaryotes. Nature 468:60–6.

